# VPS13A and VPS13C influence lipid droplet abundance

**DOI:** 10.1101/2022.06.21.497109

**Authors:** Shuliang Chen, Melissa A. Roberts, Chun-Yuan Chen, Sebastian Markmiller, Hong-Guang Wei, Gene W. Yeo, James G Granneman, James A. Olzmann, Susan Ferro-Novick

**Affiliations:** Department of Cellular and Molecular Medicine, University of California San Diego, La Jolla, CA 92093, USA; Department of Molecular and Cell Biology, University of California, Berkeley, Berkeley, CA 94720, USA; Department of Nutritional Sciences and Toxicology, University of California, Berkeley, Berkeley, CA 94720, USA; Center for Integrative Metabolic and Endocrine Research, Wayne State University School of Medicine, Detroit, MI, 48201, USA; Chan Zuckerberg Biohub, San Francisco, CA 94158, USA

**Author notes:** Address correspondence to: Susan Ferro-Novick;, James A. Olzmann. Equal contribution (Shuliang Chen, Melissa A. Roberts).

## Abstract

Lipid transfer proteins mediate the exchange of lipids between closely apposed membranes at organelle contact sites and play key roles in lipid metabolism, membrane homeostasis, and cellular signaling. A recently discovered novel family of lipid transfer proteins, which includes the VPS13 proteins (VPS13A-D), adopt a rod-like bridge conformation with an extended hydrophobic groove that enables the bulk transfer of membrane lipids for membrane growth. Loss of function mutations in VPS13A and VPS13C cause chorea acanthocytosis and Parkinson’s disease, respectively. VPS13A and VPS13C localize to multiple organelle contact sites, including endoplasmic reticulum (ER) – lipid droplet (LD) contact sites, but the functional roles of these proteins in LD regulation remains mostly unexplored. Here, we employ CRISPR-Cas9 genome editing to generate VPS13A and VPS13C knockout cell lines in U-2 OS cells via deletion of exon 2 and introduction of an early frameshift. Analysis of LD content in these cell lines revealed that loss of either VPS13A or VPS13C results in reduced LD abundance under oleate-stimulated conditions. These data implicate VPS13A and VPS13C in LD regulation and raise the intriguing possibility that VPS13A and VPS13C-mediated lipid transfer facilitates LD biogenesis.

## INTRODUCTION

Members of the highly conserved VPS13 protein family are found at organelle contact sites (Kumar et al., 2018; Leonzino et al., 2021). The founding member of the family, yeast Vps13, was first identified in *Saccharomyces cerevisiae* in a screen for genes required for <u>v</u>acuolar <u>p</u>rotein <u>s</u>orting (VPS) from the Golgi to the vacuole (Bankaitis et al., 1986). Subsequent studies revealed that Vps13 is also required for sporulation, mitochondrial homeostasis and endoplasmic reticulum (ER) autophagy (Chen et al., 2020; Park et al., 2016; Park and Neiman, 2012). Consistent with its pleiotropic roles, Vps13 localizes to the sporulation membrane and multiple ER-organelle contact sites that include the mitochondria, vacuole and endosome/vacuole (Chen et al., 2020; Park et al., 2016; Park and Neiman, 2012). While the precise function of Vps13 at these contact sites is unclear, recent studies have revealed that Vps13 is a lipid transporter that delivers glycerol lipids across membranes in vitro (Kumar et al., 2018; Li et al., 2020). Yeast encodes one *VPS13* gene, while mammals contain four genes: VPS13A, VPS13B, VPS13C and VPS13D. The yeast gene is most closely related to mammalian VPS13A and VPS13C. Loss of function mutations in VPS13A have been linked to chorea acanthocytosis, a neurological disorder that leads to Huntington-like muscle degeneration and abnormally shaped red blood cells (Rampoldi et al., 2001; Ueno et al., 2001), while mutations in VPS13C are associated with early onset Parkinson’s disease (Lesage et al., 2016). VPS13A is found at ER-mitochondria contact sites, whereas VPS13C localizes to ER-late endosome/lysosome contact sites (Kumar et al., 2018). VPS13A and VPS13C are also present at ER-lipid droplet (LD) contact sites (Kumar et al., 2018) and VPS13C is detected in high-confidence LD proteomes (Bersuker et al., 2018). Furthermore, VPS13C can be found in a distinct subdomain of LDs in mouse brown adipocyte tissue (BAT) cells (Ramseyer et al., 2018).

LDs are conserved ER-derived organelles that store excess fatty acids (FAs) in the form of neutral lipids such as triacylglycerol and cholesterol esters (Olzmann and Carvalho, 2019). Structurally, LDs consist of a central core of neutral lipids that is encircled by a phospholipid monolayer. Integral and peripheral proteins associate with the bounding phospholipid monolayer and regulate LD growth and turnover, as well as interactions with other cellular organelles (Olarte et al., 2022; Roberts and Olzmann, 2020). LDs are hubs of lipid metabolism. The size and number of LDs, as well as the mobilization of neutral lipids from LDs, are highly regulated to meet the cellular demand for energy conversion, the production of lipid signaling molecules, and the biosynthesis of phospholipids for membrane expansion (Olzmann and Carvalho, 2019). LD biogenesis begins with the deposition of neutral lipids within the ER bilayer (Thiam and Ikonen, 2021). The neutral lipids phase separate to form a lens-like structure in the ER membrane, and the LD subsequently buds from the outer leaflet of the ER into the cytosol through a process that is promoted by an LD assembly complex composed of seipin and LDAF1 (lipid droplet assembly factor 1) (Thiam and Ikonen, 2021). Interestingly, in addition to the ER, LDs make contacts with other organelles such as the mitochondria, lysosomes, and peroxisomes (Olzmann and Carvalho, 2019). Previous studies indicate a key role for contact sites in the interorganellar exchange and transfer of lipids (Prinz et al., 2020).

Because the phospholipid transfer proteins VPS13A and VPS13C are found at ER-LD contact sites (Kumar et al., 2018) and LD growth requires new phospholipids for directional emergence into the cytosol (Chorlay et al., 2019), in the current study we examined whether VPS13A and VPS13C regulate LD formation and size.

## RESULTS AND DISCUSSION

VPS13A and VPS13C are large genes that contain 71 and 83 exons, respectively. To generate knockout (KO) cell lines, we used CRISPR-Cas9 to delete exon 2 in both genes. We introduced two double-stranded breaks in VPS13A and VPS13C, which generated a frame shift downstream of exon 1 (**Figure 1**). The deletion of exon 2 was confirmed using two primer sets, including flanking primers and an exon 2 internal reverse primer (**Figure 1 and 2A**). Deletion of exon 2 resulted in the expected shift in amplicon size using the flanking primers. We observed a reduction of ∼330bp for the VPS13A amplicon and a reduction of ∼464bp for the VPS13C amplicon. No amplicon was detected when using exon 2 internal reverse primers (**Figure 2A**). Furthermore, loss of the VPS13A and VPSC13C proteins was confirmed by immunoblotting (**Figure 2B**). These data confirm the successful deletion of exon 2 in VPS13A and VPS13C and demonstrate that the frameshift we generated disrupts protein expression.

**Figure 1.**
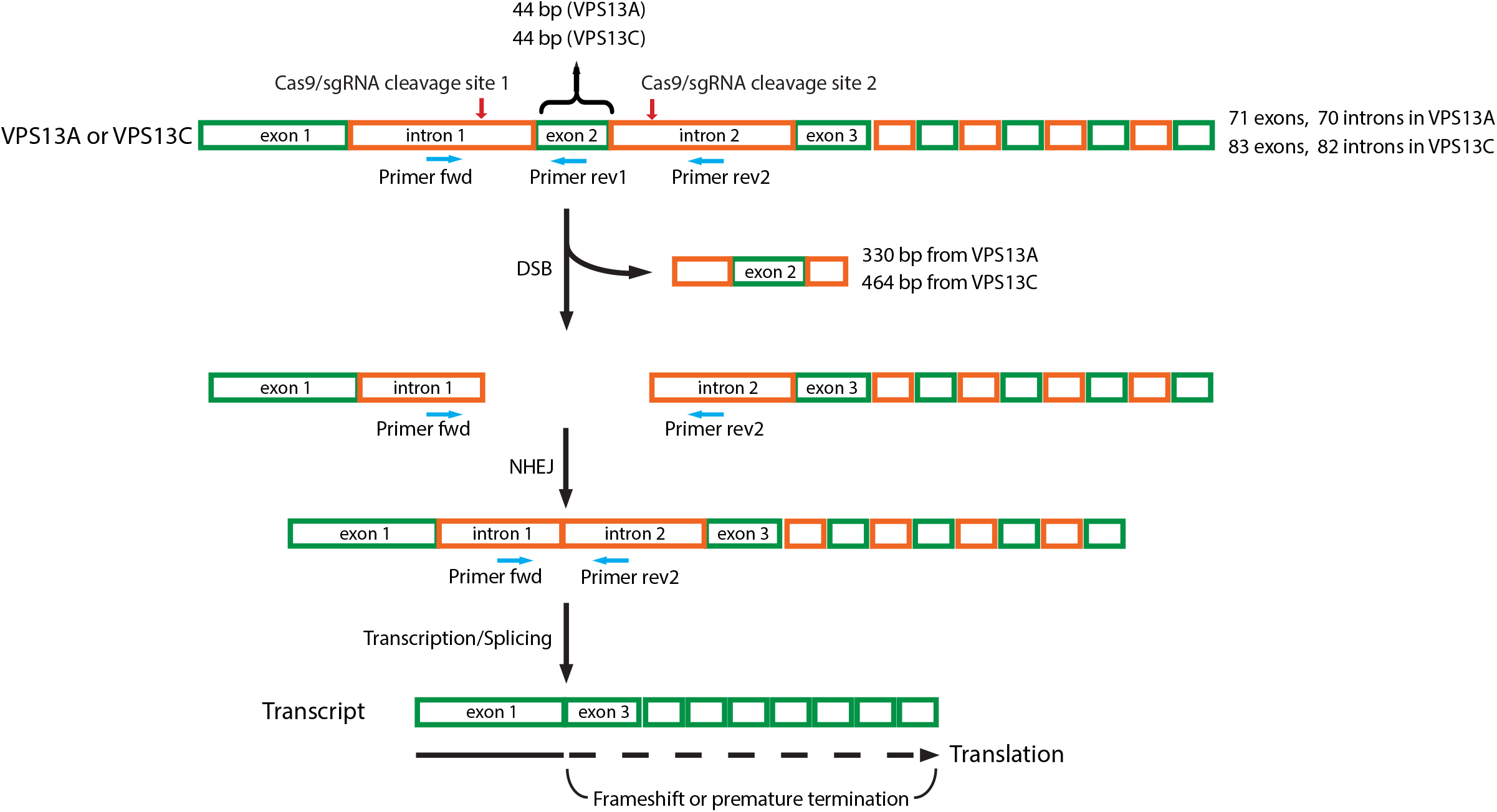
Schematic diagram of the strategy used to knock-out VPS13A and VPS13C using the CRISPR/Cas9 system. Two sgRNAs were designed to delete exon 2 of human VPS13A or VPS13C. Exon 2 in the VPS13A and VPS13C genes is 44 bp in length. One sgRNA targeted intron 1 on the 5′ sides of exon 2, the other targeted intron 2 on the 3′ side of the exon 2. Cas9/sgRNAs create two double-stranded breaks (DSBs) that excise the exon 2 containing DNA fragment, 330 bp from VPS13A and 464 bp from VPS13C. Cells repair DSBs via non-homologous end joining (NHEJ) mediated re-ligation of broken DNA ends, but exon 2 was missing in the repaired VPS13A and VPS13C genes and their transcripts. Since the size of exon 2 is 44 bp, the transcript contains frameshift mutations downstream of exon 1, leading to the incorporation of incorrect amino acids in the proteins or premature termination during translation. The primer pair of Primer-fwd (binds to intron 1) and Primer-rev1 (binds to exon 2) was used to detect the deletion of exon 2, and Primer-fwd (binds to intron 1) and Primer-rev2 (binds to intron 2) was used to detect the occurrence of NHEJ between intron 1 and intron 2. Green boxes indicate exons, and orange boxes indicate introns. Red arrows point to the Cas9/sgRNA cleavage site. Blue arrows indicate primers for PCR-based validation.

**Figure 2.**
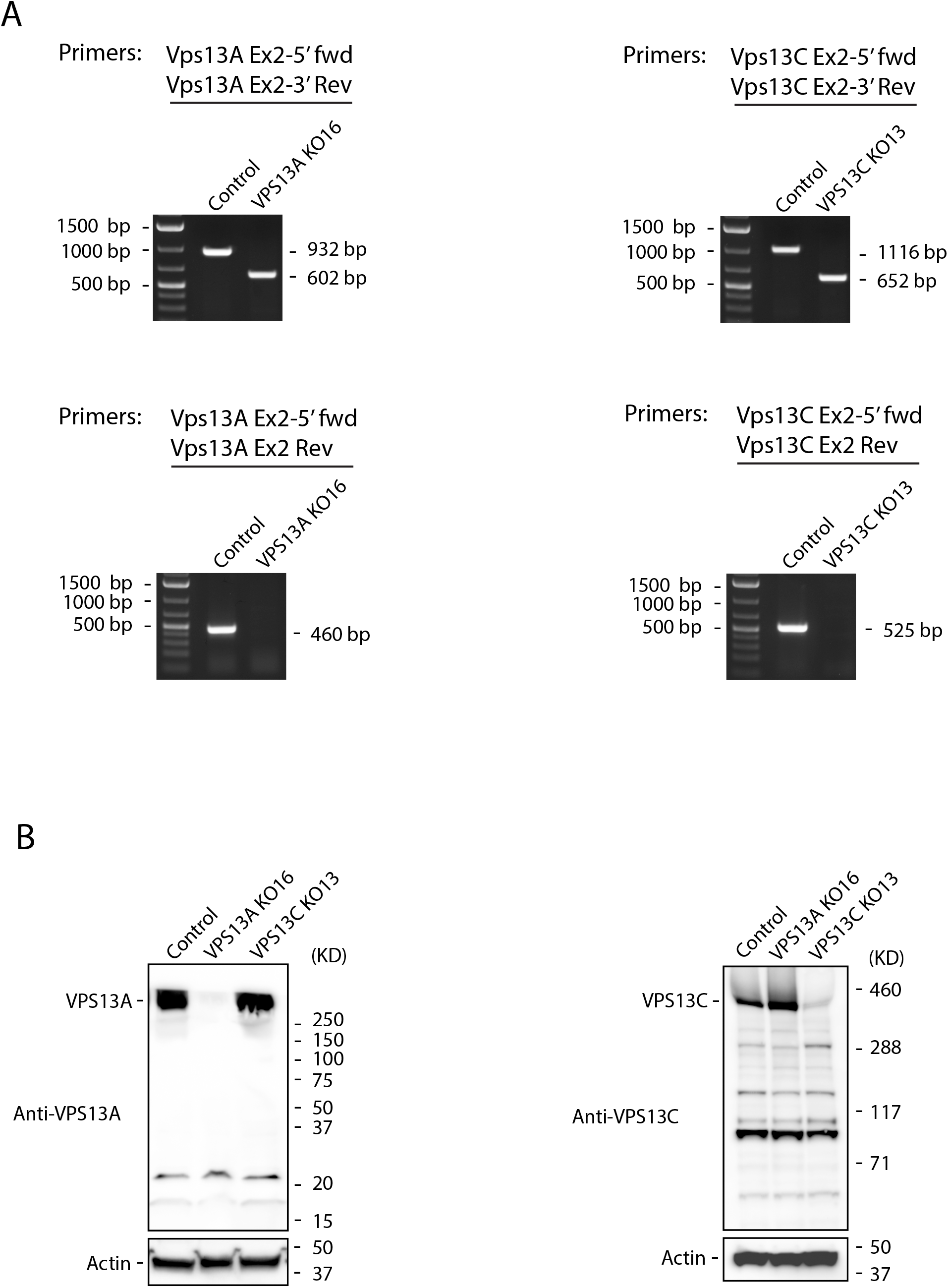
Validation of the VPS13A and VPS13C knock-outs. (A) PCR-based validation. Left panel, genomic DNA extracted from the VPS13A knock-out cell line (VPS13A KO16) was analyzed for the deletion of exon 2. A 932 bp PCR product was amplified in control cells by a set of primers, Vps13A Ex2-5’ fwd, that bind to intron 1 and Vps13A Ex2-3’ Rev, that bind to intron 2. By contrast, the PCR product from VPS13A KO16 was 602 bp, reflecting a deletion of the exon 2 containing DNA fragment. Primers Vps13A Ex2-5’ fwd and Vps13A Ex2 Rev were used to directly detect exon 2. A predicted 460 bp band was visualized in control cells, while the band was not detectable in VPS13A KO16. Right panel, genomic DNA extracted from the VPS13C knock-out cell line (VPS13C KO13) was analyzed for the deletion of exon 2. In VPS13C KO13 cells, primers Vps13C Ex2-5’ fwd, that bind to intron 1, and Vps13C Ex2-3’ Rev, that bind to intron 2, amplified a band of 652 bp. This band was shorter than the band from control cells (1116 bp) as a consequence of the deletion of the exon 2 containing DNA fragment. Using primers Vps13C Ex2-5’ fwd, that bind to intron 1, and Vps13C Ex2 Rev, that bind to exon 2, a 525 bp amplicon from control cells was produced. Because of the absence of exon 2, the amplicon was not produced from VPS13C KO13 cells. (B) Western blot analysis of the knock-out cells. Cell lysates from control U-2 OS, VPS13A KO16 and VPS13C KO13 cells were immunoblotted with anti-VPS13A antibody (left panel) and anti-VPS13C antibody (right panel). Actin was used as a loading control.

To examine a potential role for VPS13A and VPS13C in regulating LDs, we quantified the number and size of LDs in wild type, VPS13A KO, and VPS13C KO U-2 OS cells. Because U-2 OS cells have very few detectable LDs under basal conditions, we stimulated LD biogenesis by treating cells for 24 hours with 200 *µ*M oleate complexed with bovine serum albumin (BSA). Quantification of LDs from each knock-out (>1200 cells) revealed a modest but significant reduction in the number of LDs per cell in both the VPS13A KOs and VPS13C KOs (**Figure 3A and B**). The defect was, however, more dramatic in the VPS13C KOs (**Figure 3B**), and there was no significant change in the distribution of LD size in either KO cell line (**Figure 3C**). These data indicate that loss of VPS13A and VPS13C leads to reduced LD abundance. Through their chorein domain, VPS13A and VPS13C act as phospholipid transfer proteins at organelle contact sites (Leonzino et al., 2021). Our results indicate that both VPS13A and VPS13C are required to form wild type-levels of LDs during oleate supplementation in U-2 OS cells.

**Figure 3.**
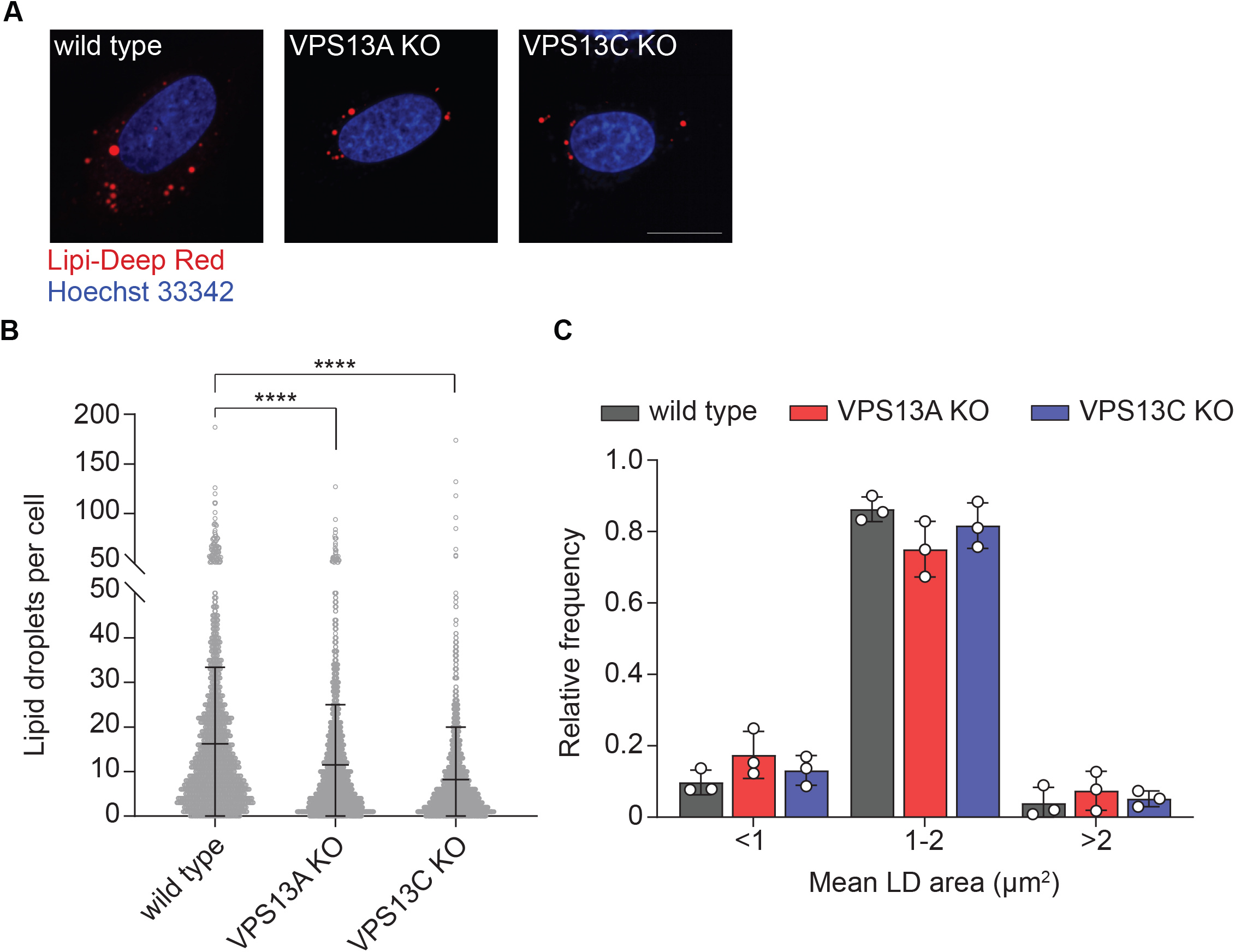
Lipid droplet abundance is reduced in VPS13A or VPS13C KO cells. **A)** Control, VPS13A KO, and VPS13C KO U-2 OS cells treated with 200 *µ*M oleate for 24 hr were stained with Lipi-Deep red (LDs) and Hoechst 33342 (nuclei) and visualized using confocal microscopy. Images are representative of at least 1200 cells imaged for each cell line. Scale bar, 20 *µ*M. LD number and area per cell are quantified in panels **B** and **C**, respectively. Data represent the mean ± standard deviation of three biological replicates. \*\*\*\**p* < 0.0001 by unpaired, two-tailed *t-*test.

Phospholipids encircle the neutral lipid core of LDs, acting as surfactants that prevent aberrant LD fusion (Guo et al., 2008; Krahmer et al., 2011) and enabling the directional emergence of lipid droplets into the cytosol during biogenesis (Chorlay et al., 2019). It is possible that VPS13A and VPS13C transfer phospholipids to LDs to regulate LD fusion and/or biogenesis. Under the conditions examined, loss of VPS13A and VPS13C did not impact LD size, suggesting that VPS13A and VPS13C do not affect fusion. We propose that the contribution of VPS13A and VPS13C to LD regulation may differ depending on the cell line or tissue examined. For example, VPS13C depletion in cultured murine BAT results in reduced LD size due to an increase in ATGL-mediated lipolysis (Ramseyer et al., 2018). Conversely, VPS13A depletion in MRC-5 lung fibroblasts increased the number of LDs through an as yet undetermined mechanism (Yeshaw et al., 2019). Our data implicate VPS13A and VPS13C in LD regulation in U-2 OS cells and highlights the importance for future studies to understand the mechanistic underpinnings of these genes in diverse cell types under different metabolic conditions.

## MATERIALS AND METHODS

### Cell culture and oleate treatment

U-2 OS cells were cultured in DMEM containing 4.5 g/L glucose and L-glutamine (Corning) supplemented with 10% fetal bovine serum (FBS, Gemini Bio Products) at 37°C with 5% CO2. The day before oleate treatment, cells were seeded at 400,000 cells per well into 24-well glass bottom plates coated with poly-L-lysine. The next day, cells were treated with 200 *µ*M oleate-BSA complex for 24 hr prior to imaging. For the oleate treatment, the following reagents were combined to obtain 1 ml of oleate-BSA complex: 890 *µ*l DMEM, 100 *µ*l BSA in PBS (100x, Sigma #A8806), and 10 *µ*l oleate stock (200 mM stock in ethanol, Sigma #O1383). The mixture was vortexed and added to cells at a 1:10 v/v ratio (final concentration, 200 *µ*M oleate).

### Generation of VPS13A and VPS13C knock-out cell lines

Human VPS13A and VPS13C knock-outs in U-2 OS cell lines were generated by two double-stranded breaks (DSBs) using CRISPR/Cas9 approaches. Single-guide RNAs (sgRNA) targeting introns at the 5′ and 3′ sides of exon 2 of VPS13A or VPS13C (Figure 1) were designed using the CHOPCHOP web tool (Labun et al., 2021). The sgRNA oligonucleotides listed in Table 1 were synthesized (IDT), annealed, and ligated into the BbsI site of pSpCas9(BB)-2A-Puro vector (PX459, Addgene #62988). U-2 OS cells were transfected with two PX459 encoding sgRNAs using lipofectamine 2000 (Invitrogen) following the manufacturer’s instructions. One sgRNA targeted intron 1 at the 5′ side of exon 2, and the other sgRNA targeted intron 2 at the 3′ side of exon 2. 24 hours post-transfection, cells were selected by treatment with 2 *µ*g/ml puromycin for 72 h. Single puromycin-resistant colonies were established by subcloning populations into 96-well plates. Genomic DNA was isolated from the expanded individual colonies and used for screening the exon 2 deletion in VPS13A or VPS13C by PCR.

**Table 1.**
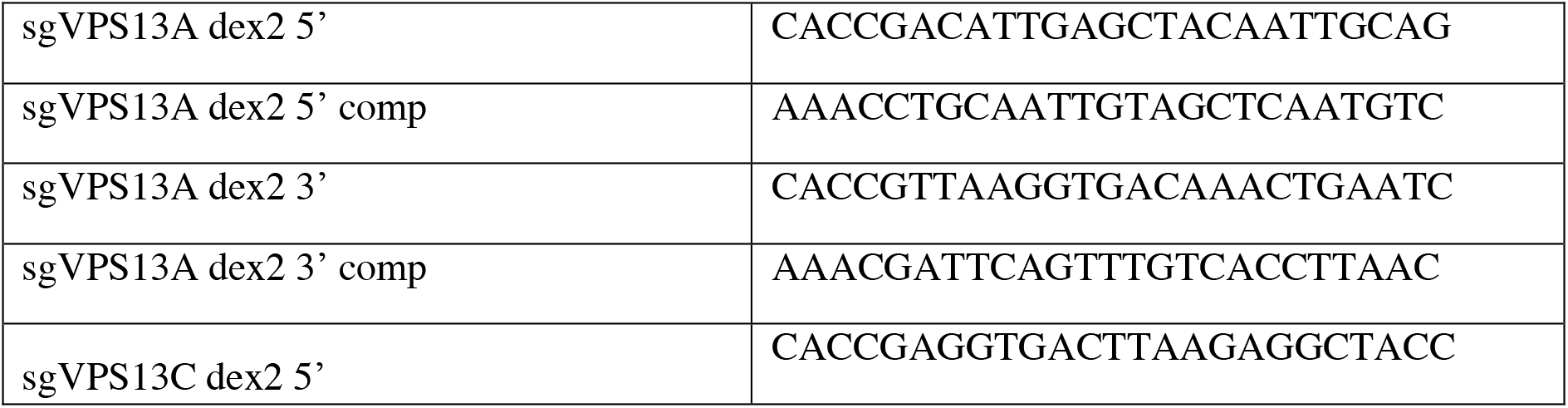
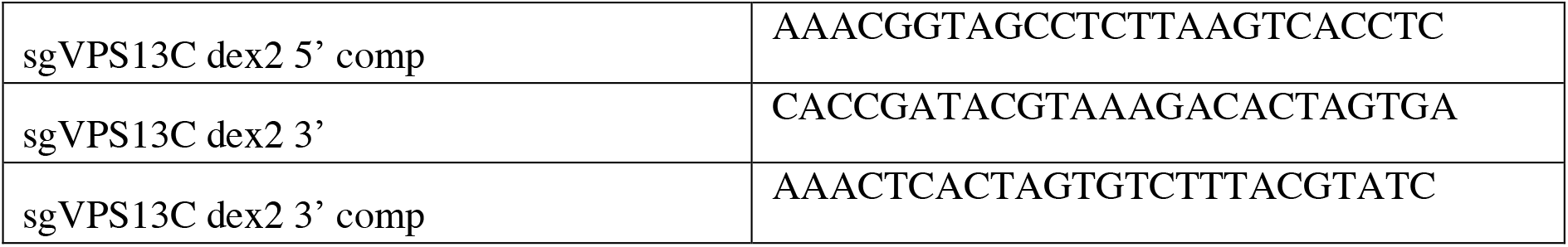
Oligonucleotides used in this study to construct sgRNA-encoding PX459 vectors for VPS13A and VPS13C knock-outs.

### PCR-based validation of the VPS13A and VPS13C knock-out cell lines

Puromycin-resistant cell clones were seeded in 96-well plates and cultured in DMEM medium. When cells reached 80-100% confluency, the medium was aspirated and cells were washed twice with PBS. To extract genomic DNA, the cells were re-suspended in 20 *μ*l of DNA Extraction Solution (Lucigen), and incubated at 68°C for 6 min, followed by a 98°C incubation for 2 min. The PCR validation system was set up using GoTaq Green Master Mix (Promega) with the following components: 7.5 *μ*l of GoTaq Green Master Mix; 1 *μ*l of genomic DNA extract; 2 *μ*l of forward primer (2 *μ*M); 2 *μ*l of reverse primer (2 *μ*M); and H_2_O up to 15 *μ*l. Primers for PCR-based validation of VPS13A and VPS13C knock-outs are listed in Table 2. PCR was performed in a thermocycler using the following cycling conditions: 98°C for 10 min; 35 cycles for the extending reaction (98°C for 45 sec, 55°C for 45 sec, 72°C for 2 min), and 72°C for 10 min. PCR products were analyzed by 1% agarose gel electrophoresis.

**Table 2.**
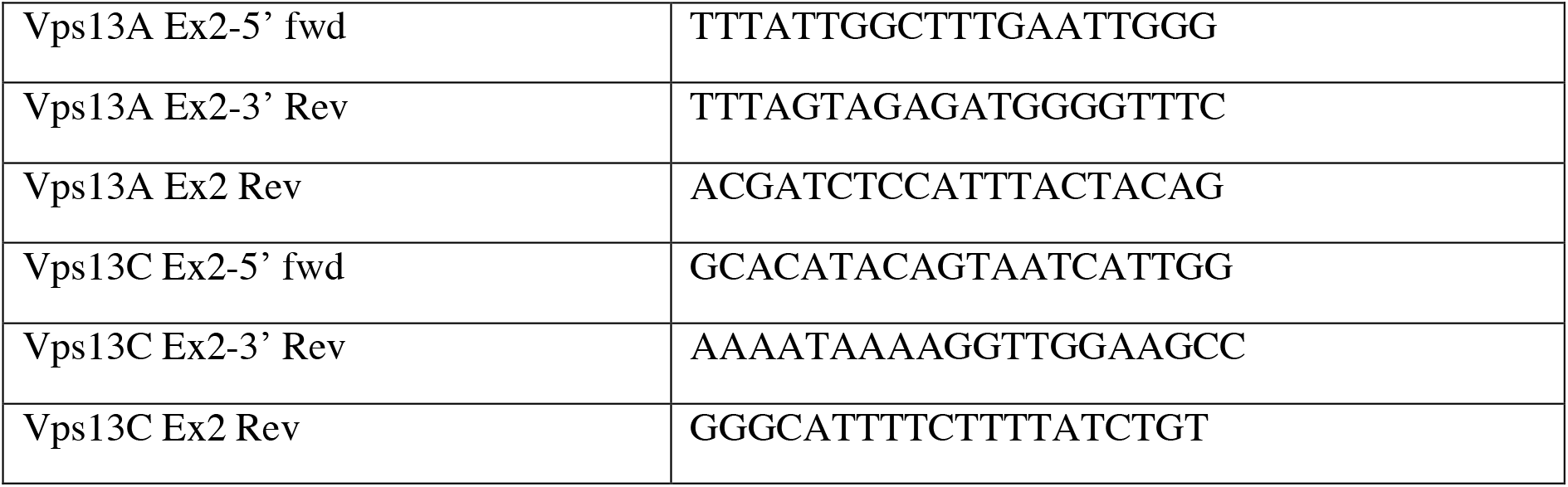
Primers for PCR-based validation of VPS13A and VPS13C knock-outs.

### Western blot analysis

To ultimately confirm the VPS13A and VPS13C knock-outs, western blot analysis was performed. Cells with exon 2 deletion were lysed in RIPA buffer and the presence of VPS13A and VPS13C was detected using anti-VPS13A (Sigma, HPA021662) and anti-VPS13C (Ramseyer et al., 2018) antibodies, respectively. Cell clones with undetectable VPS13A (VPS13A KO16) and VPS13C (VPS13C KO13) were used for further analysis.

### Confocal microscopy

U-2 OS cells were grown in 24-well glass bottom plates (170 *µ*m coverglass bottom; Eppendorf #0030741021) coated with poly-l-lysine and treated with 200 *µ*M oleate-BSA complex for 24 hr. Cells were incubated with 0.5 *µ*M Lipi-Deep Red neutral lipid stain (Dojindo #LD04-10) for 2 hr and 5 *µ*g/mL Hoeschst 33342 nucleic acid stain (Invitrogen #H3570) for 30 min at 37°C. Cells were then washed twice with PBS and imaged in fresh medium lacking phenol red.

Live cells were imaged using an Opera Phenix Plus High-Content Screening System (Perkin Elmer) confocal microscope equipped with a 63X water immersion objective using DAPI and Cy-5 filters. Cells were imaged at 37 °C with 5% CO2. Z-stacks of 0.5-μm slices totaling 8 μm in thickness were acquired. Images were merged and brightness and contrast adjusted using Fiji (https://imagej.net/software/fiji/).

### Lipid droplet quantification

LDs were quantified by creating a custom analysis sequence using Harmony (version 4.9) high-content analysis software. For each field, maximum projection Z-stacks were processed with advanced flatfield correction. Nuclei and cytoplasm were defined using the DAPI and Cy-5 channels, respectively, and border cells were automatically excluded from analyses. LDs were defined using the “Find Spots” task (Lipi-Deep red stain, Cy-5 channel) thresholding for size, intensity, and roundness. For each cell, the average lipid droplet number and area (*µ*m^2^) were quantified. LD quantification data from three biological replicates were graphed and analyzed in Prism 9 (GraphPad). For each cell line, at least 1200 cells were analyzed. *P*-values for pairwise comparisons were calculated via unpaired, two-tailed *t*-test.

## ACKNOWLEDGEMENTS

This research was supported by grants from the National Institutes of Health (R01GM112948 and R01DK128099 to J.A.O., R01DK076629 to J.G.G., R35GM131681 and R01NS117440 to S.F-N.). This work was partially supported by National Institutes of Health grants HG004659 and HG009889 to G.W.Y. G.W.Y is also supported by an Allen Distinguished Investigator Award, a Paul G. Allen Frontiers Group advised grant of the Paul G. Allen Family Foundation.

